# Age-related susceptibility of ferrets to SARS-CoV-2 infection

**DOI:** 10.1101/2021.08.16.456510

**Authors:** Mathias Martins, Maureen H.V. Fernandes, Lok R. Joshi, Diego G. Diel

## Abstract

Susceptibility to SARS-CoV-2 and the outcome of COVID-19 have been linked to underlying health conditions and the age of affected individuals. Here we assessed the effect of age on SARS-CoV-2 infection using a ferret model. For this, young (6-month-old) and aged (18-to-39-month-old) ferrets were inoculated intranasally with various doses of SARS-CoV-2. By using infectious virus shedding in respiratory secretions and seroconversion, we estimated that the infectious dose of SARS-CoV-2 in aged animals is ∼32 plaque forming units (PFU) per animal while in young animals it was estimated to be ∼100 PFU. We showed that viral replication in the upper respiratory tract and shedding in respiratory secretions is enhanced in aged ferrets when compared to young animals. Similar to observations in humans, this was associated with higher expressions levels of two key viral entry factors - ACE2 and TMPRSS2 - in the upper respiratory tract of aged ferrets.

---

In late December 2019, several cases of viral pneumonia of unknown etiology were described in a cluster of people in Wuhan, Hubei province, China. The causative virus severe acute respiratory syndrome coronavirus 2 (SARS-CoV-2) was subsequently discovered and the disease named coronavirus disease 19 (COVID-19)^1^. Genome sequence analysis determined that SARS-CoV-2 is closely related to the bat SARS-like CoV RaTG13, a virus that was identified in horseshoe bats in the same geographic region in China. Given the close genetic and phylogenetic relationship of SARS-CoV-2 and SARS-like CoV found in bats, these animals are currently considered the likely source of the ancestral virus that originated SARS-CoV-2^2–4^. Importantly, epidemiological investigations on the early clusters of COVID-19 revealed a strong link of the initial human cases with the Huanan Seafood Wholesale Market in Wuhan, where presumably SARS-CoV-2 may have been transmitted to humans through a yet unknown intermediate animal host^2, 5^. Since the description of first cases, SARS-CoV-2 spread across the world causing an unprecedented global pandemic that by August 2021 incurred in over 200 million human cases and more than 4.3 million deaths (https://covid19.who.int/).

Coronaviruses are positive sense, single stranded RNA viruses of the family *Coronaviridae*. Although coronaviruses usually cause mild respiratory infection in humans, they can also infect a range of other species resulting in broad clinical outcomes, varying from subclinical or mild-to-severe respiratory- to gastroenteric infections, which in some cases can lead to fatal disease^6^. Historically, several zoonotic coronaviruses have jumped into humans, including severe acute respiratory syndrome coronavirus (SARS-CoV), which was described in China in 2002, and Middle East respiratory syndrome coronavirus (MERS-CoV) identified in Saudi Arabia in 2012. Interestingly, SARS-CoV, MERS-CoV and SARS-CoV-2 present remarkable differences in infection outcomes. The fatality rates of SARS-CoV and MERS-CoV, for example, are much higher (∼10% and ∼35%, respectively) than that described for SARS-CoV-2 (<3%)^7, 8^. The clinical outcomes of SARS-CoV-2 infection are variable, ranging from asymptomatic infections - which represent most cases - to multiple organ failure and death^9^. Importantly, the most severe cases of COVID-19 resulting in death have been associated with other underlying health conditions^10, 11^. In addition, age is another risk factor that has been associated with distinct and often more severe outcomes of SARS-CoV-2 infection. While most human infections with SARS-CoV-2 lead to subclinical disease or only mild symptoms in young healthy adults, disease severity and mortality rates increase with age among individuals older than 30 years^12, 13^. Additionally, viral load, which is a direct measure of virus replication and that has been linked to severe disease outcomes, has also been shown to increase with the age of affected individuals^12, 14^. The mechanisms underlying the different SARS-CoV-2 infection outcomes and the contribution of comorbidities or age to these diverse outcomes, however, remains unknown.

The first step of SARS-CoV-2 infection involves binding of the viral spike (S) protein, more precisely the S receptor binding domain (RBD) to angiotensin-converting enzyme 2 (ACE2)– the viral cognate receptor^15, 16^. Following S RBD-ACE2 binding, the S protein is cleaved by host proteases (e.g. furin) into the S1/S2 subunits. The next step that takes place is cleavage of the S2 subunit at the S2’ site by the transmembrane serine protease 2 (TMPRSS2), which leads to conformational changes that expose the fusion peptide enabling membrane fusion and completion of viral entry into host cells^16^. In humans, ACE2 is expressed in various cells and tissues, with high levels of the protein being expressed in the upper respiratory tract (URT)^17–19^. Notably, differential ACE2 and TMPRSS2 expression have been described in young and old people, with a higher percentage of ACE2 and TMPRSS2 expressing cells being detected in the nasal brushing of older people when compared to young individuals^20, 21^.

Here we assessed the effect of age on SARS-CoV-2 infection in ferrets. We compared the susceptibility of young (6-month-old) and aged (18- to 39-month-old) ferrets to four different doses of SARS-CoV-2. Viral replication, viral load and shedding in respiratory secretions and feces were monitored by rRT-PCR, virus isolation and titrations for 14 days post-inoculation (pi), while seroconversion was assessed by virus neutralization assays. The virological and serological findings were used to estimate the median infectious dose (ID_50_) of SARS-CoV-2 in young and aged ferrets. Additionally, expression of ACE2 and TMPRSS2 were assessed in upper and lower respiratory tract of ferrets and correlated with outcomes of SARS-CoV-2 infection and replication.

## Results

### Clinical parameters following intranasal inoculation of SARS-CoV-2 in young and aged ferrets

To assess the susceptibility of ferrets to SARS-CoV-2, a total of 40 (20 young [6-month-old] and 20 aged [18- to 39-month-old]) animals were allocated in 10 experimental groups (n = 4 per group). Animals were mock-inoculated (control group) or inoculated intranasally with different doses of SARS-CoV-2 (10^1^, 10^2^, 10^3^, and 10^6^; Fig. 1a). Virus inoculated ferrets were housed in the Animal Biosafety Level 3 (ABSL-3) facility at the East Campus Research Facility (ECRF) at Cornell University. While control animals (four young and four aged animals) were kept under ABLS-1 conditions. Both inoculated and control ferrets were housed individually in Horsfall HEPA-filter cages throughout the 14-day experimental period. Following inoculation, clinical parameters, including temperature, body weight, activity, and signs of respiratory disease were monitored daily. The body temperature in both age groups remained within physiological ranges throughout the experimental period, and no differences were observed between experimental groups (Fig. 1b, c). The body weight had slight variation in all groups, but no difference was noticed between SARS-CoV-2-inculated groups to the mock-control ferrets (normalized, day 0 represent 100%), in both young and aged animals (Fig. 1d, e). No marked differences in daily activity of mock-control- or SARS-CoV-2-inoculated animals was noticed. It is important to note that, young and aged ferrets naturally exhibit differences in behavior^22^. While young animals are more active and curious, aged ferrets are less active and they are more restful and calmer than young animals, and this was also observed in our study. Additionally, no clinical signs of respiratory disease were observed in any of the SARS-CoV-2-inoculated or control mock-inoculated ferrets throughout the experimental period.

**Fig. 1.**
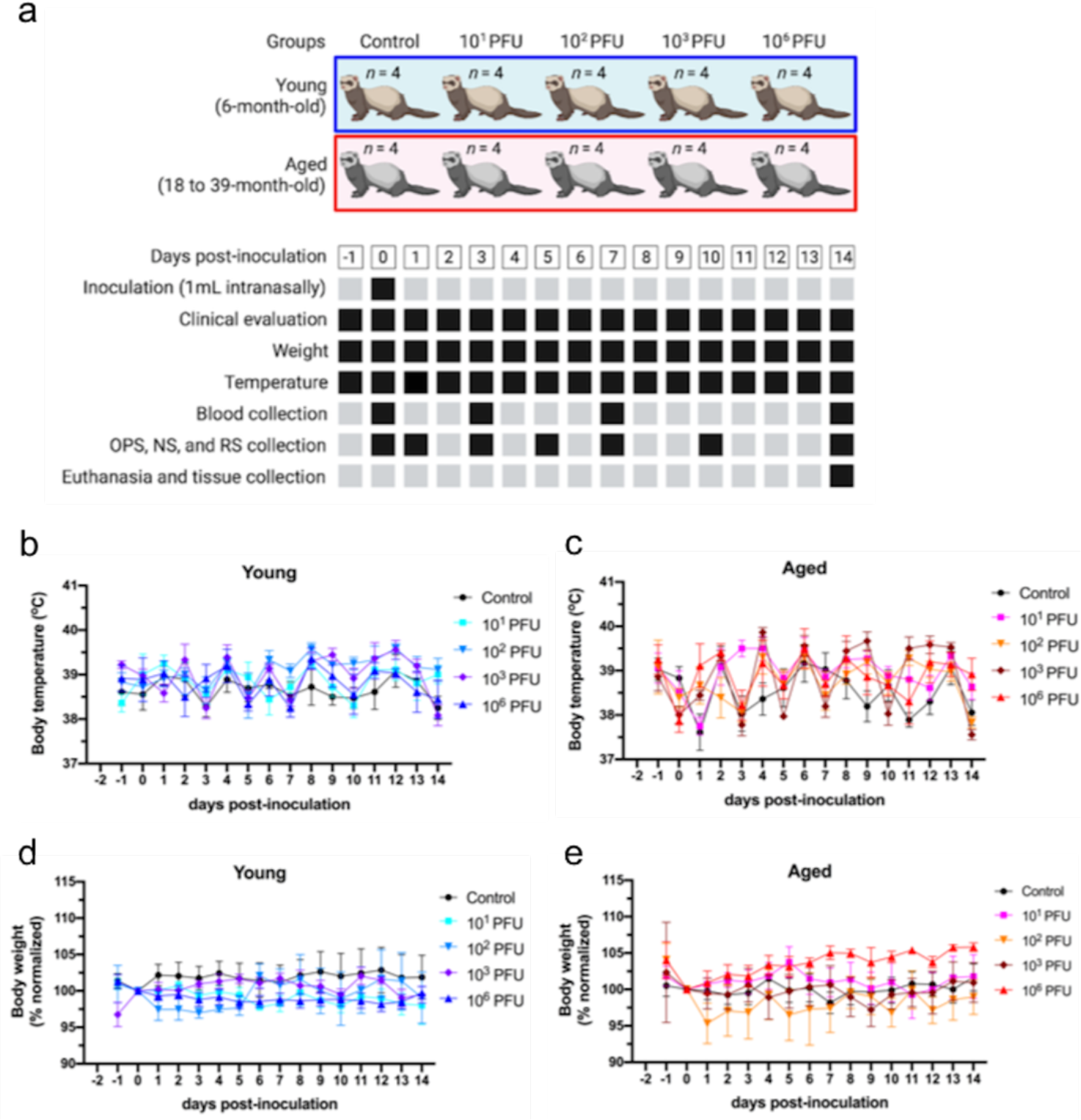
Experimental design, body weight and temperature following SARS-CoV-2-inoculation. Forty ferrets (*Mustela putorius furo*) - Twenty young (6-month-old) and twenty aged (18 to 39-month-old [30.5±5.3 - average±SD]) were allocated to ten experimental groups (4 animals per group). Animals were inoculated intranasally with 1 ml (0.5 ml per nostril) MEM (mock control groups) or with a 1 ml virus suspension containing 10^1^, 10^2^, 10^3^ and 10^6^ PFU of SARS-CoV-2 isolate NY67-20 on day 0. All animals were maintained individually in Horsfall HEPA-filtered cages. Clinical parameters, including temperature, body weight, activity, and signs of respiratory disease were monitored daily for 14 days post-inoculation (pi). Oropharyngeal- (OPS), nasal- (NS), rectal swab (RS), and blood were collected on various time points pi. Animals were humanely euthanized on day 14 pi. Following necropsy, tissues were collected and processed for rRT-PCR and virus isolation. Black hatched squares represent actual collection/measure time points for each sample type/parameter described in the figure (**a**). Body temperature following intranasal SARS-CoV-2 inoculation recorded throughout the experimental period in young (**b**) or aged ferrets (**c**). Body weight measurements throughout the experimental period in young (**d**) or aged ferrets (**e**) expressed as per cent from weight on day 0 (individual ferret weight was normalized to day 0, which represents 100%).

### Virus shedding in respiratory secretions and feces following SARS-CoV-2 inoculation

The dynamics of SARS-CoV-2 replication and shedding were monitored in respiratory secretions and feces by rRT-PCR following inoculation. Oropharyngeal- (OPS), nasal- (NS) and rectal swab (RS) samples were collected on days 0, 1, 3, 5, 7, 10 and 14 post-inoculation (pi) (Fig. 1a). Viral RNA was detected throughout the experiment in young and aged ferrets in SARS-CoV-2-inoculated animals in varying levels until day 14 pi. Higher levels of viral RNA were detected in OPS when compared to NS and RS (Fig. 2). Detection of viral RNA in animals inoculated with 10^1^ PFU of SARS-CoV-2 was restricted to a single aged ferret on day 3 pi, in which viral RNA was detected in OPS and RS samples (Fig. 2a, c). In the groups inoculated with 10^2^ PFU, 2/4 young ferrets tested positive by rRT-PCR on days 1-5 pi, while 4/4 aged animals were positive on days 1-7 pi. Importantly, viral RNA loads in OPS and RS samples were significantly higher in aged animals on days 3 (*p*<0.02), 7 and 10 pi (*p*<0.001; *p*<0.02) (Fig. 2d, f). In the groups inoculated with 10^3^ PFU, all young (4/4) and aged (4/4) ferrets tested positive for SARS-CoV-2 RNA in OPS samples between days 1-5 pi and viral RNA loads were similar between young and aged animals (Fig. 2g).

**Fig. 2.**
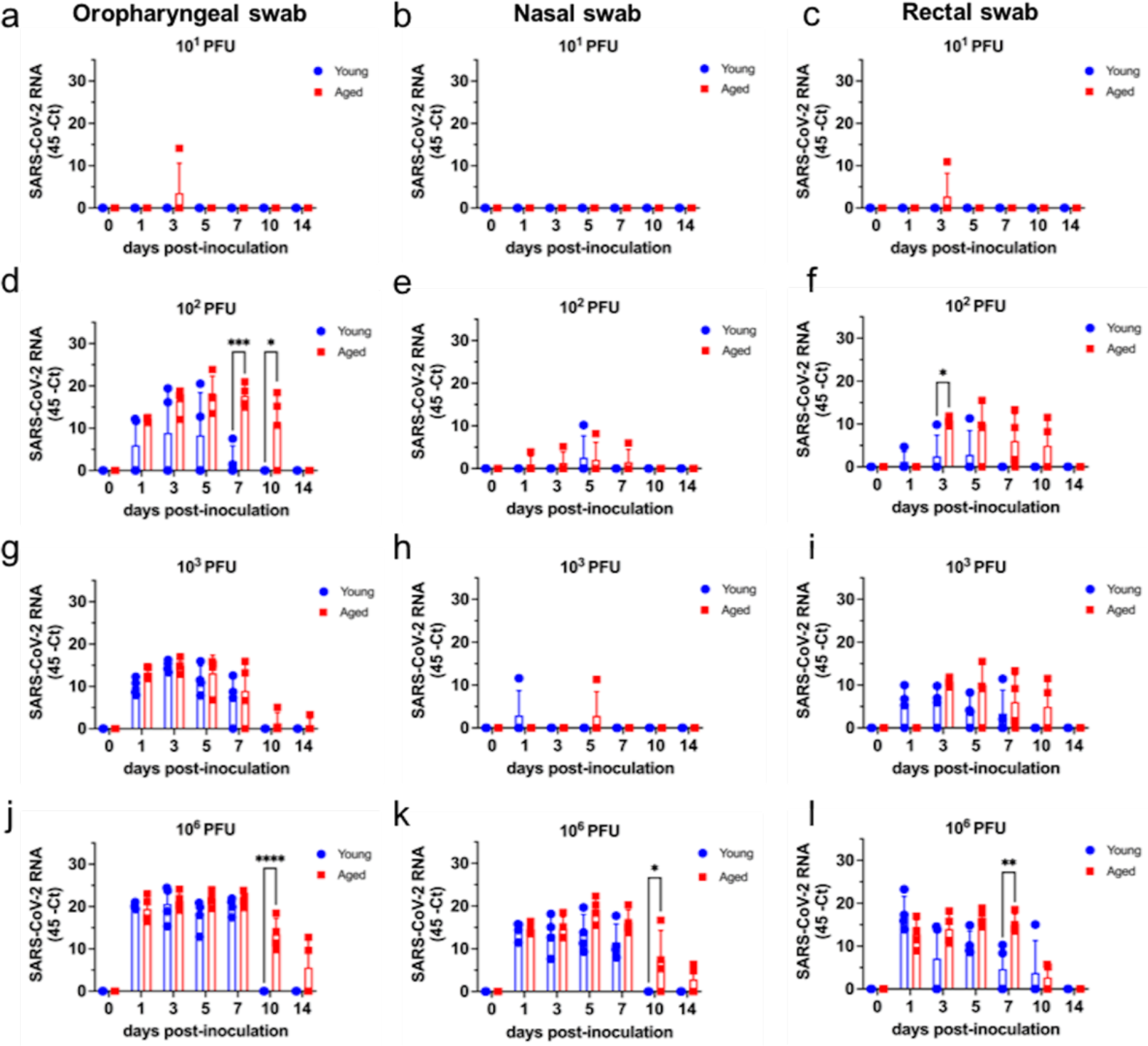
Viral RNA in respiratory secretion and feces following SARS-CoV-2 inoculation. Detection of viral RNA in Oropharyngeal- (OPS), nasal- (NS) and rectal swab (RS) samples (**a**, **b**, **c**, respectively) in young and aged ferrets inoculated with 10^1^ PFU of SARS-CoV-2. Detection of viral RNA in OPS, NS and RS samples (**d**, **e**, **f**, respectively) in young and aged ferrets inoculated with 10^2^ PFU of SARS-CoV-2. Detection of viral RNA in OPS, NS and RS samples (**g**, **h**, **i**, respectively) in young and aged ferrets inoculated with 10^3^ PFU of SARS-CoV-2. Detection of viral RNA in OPS, NS and RS samples (**j**, **k**, **l**, respectively) in young and aged ferrets inoculated with 10^6^ PFU of SARS-CoV-2. Viral RNA loads are expressed as 45 rRT-PCR cycles minus the actual Ct value. **** = *p*<0.0001; *** = *p*<0.001; ** = *p*<0.002 * = *p*<0.02.

Viral RNA was detected in RS samples in 2-3/4 animals in both young and aged groups until day 10 pi (Fig. 2i). In the groups inoculated with 10^6^ PFU, SARS-CoV-2 RNA was consistently detected in OPS, NS and RS samples, between days 1-5 pi in both young and aged animals (Fig. 2j, k, l). Interestingly, marked differences in SARS-CoV-2 RNA load were observed in NS on day 7 pi (*p*<0.02), and on OPS and RS samples on day 10 pi (*p*<0.0001; *p*<0.002, respectively) among young and aged ferrets.

To assess shedding of infectious virus in young and aged ferrets inoculated with SARS-CoV-2-, rRT-PCR positive respiratory- and fecal samples were subjected to virus isolation in cell culture. Infectious SARS-CoV-2 was isolated from OPS samples from 2/4 young ferrets and from 4/4 aged ferrets inoculated with 10^2^ PFU in at least one time point following infection (Fig. 3a, b). Higher frequency of infectious virus shedding was detected between days 3-5 pi (Fig. 3a, b). In the group inoculated with 10^3^ PFU, infectious virus shedding was detected in OPS of all 4/4 young and aged ferrets (Fig. 3c, d). No infectious virus was detected in NS or RS samples in the groups inoculated with 10^2^ or 10^3^ PFU (Fig. 3a, b, c, d). In the group inoculated with 10^6^ PFU, SARS-CoV-2 was isolated in all 4/4 young and aged ferrets, with all animals in the aged group shedding infectious virus between days 1-7 pi (Fig. 3f). Infectious SARS-CoV-2 was also isolated from NS samples from 2/4 young- and 4/4 aged animals (Fig. 3e, f).

**Fig. 3.**
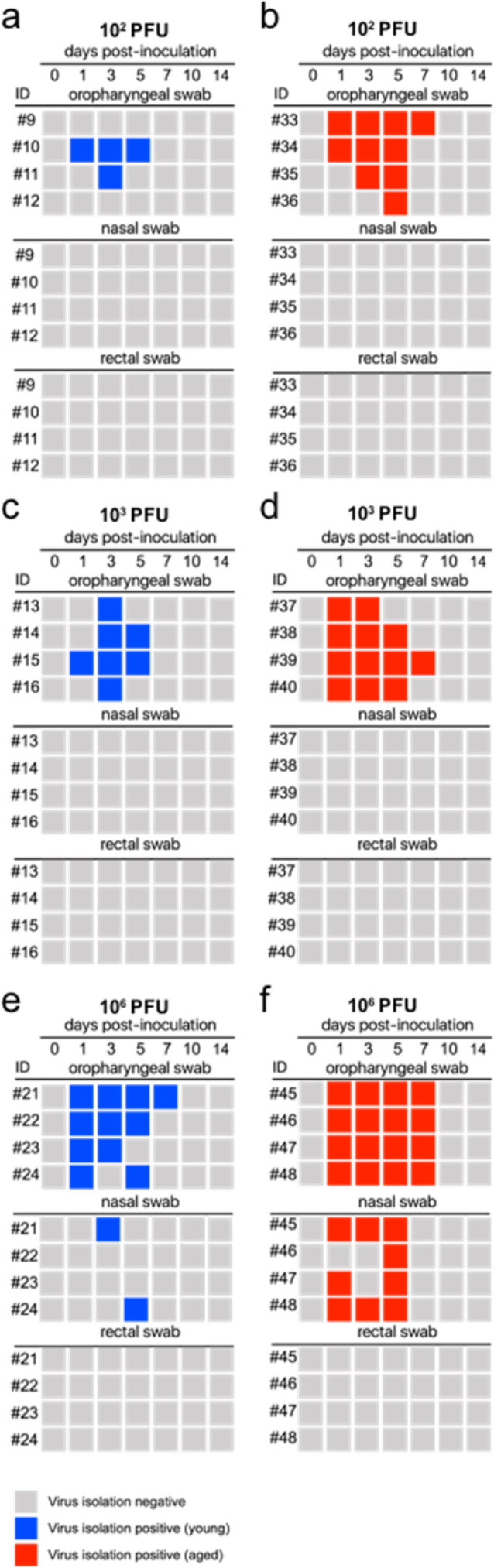
Shedding of infectious SARS-CoV-2 in respiratory secretions and feces. The rRT-PCR positive respiratory- and fecal samples were subjected to virus isolation in cell culture. Detection of infectious virus in Oropharyngeal- (OPS), nasal- (NS) and rectal swab (RS) samples of young (**a**) and aged (**b**) ferrets inoculated with 10^2^ PFU of SARS-CoV-2. Detection of infectious virus in OPS, NS and RS samples of young (**c**) and aged (**d**) ferrets inoculated with 10^3^ PFU of SARS-CoV-2. Detection of infectious virus in OPS, NS and RS samples of young (**e**) and aged (**f**) ferrets inoculated with 10^6^ PFU of SARS-CoV-2. At least three blind passages were conducted with each sample. Isolation of SARS-CoV-2 was confirmed by immunofluorescence assay using SARS-CoV-2-specific antibodies.

In addition to virus isolation, all the OPS samples that tested positive by rRT-PCR were subjected to virus quantitation. Peak viral titers were detected between days 3-5 pi (Fig. 4). In general, viral titers and the frequency of animals shedding detectable infectious SARS-CoV-2 were higher in the aged group, when compared to young animals (Fig. 4). In animals inoculated with 10^2^ PFU, statistically significant differences in virus shedding between young and aged ferrets was observed on day 5 pi (*p*<0.002) (Fig. 4a). While 4/4 aged ferrets shed virus (titers ranging from 1.0 to 2.8 log10TCID_50_.mL^-1^ [50% tissue culture infectious dose per milliliter]), only 1/4 young ferret shed 1.0 log10TCID_50_.mL^-1^ (Fig. 4a). In animals inoculated with 10^3^ PFU, all 4/4 young and aged animals shed infectious virus (Fig. 3c, d) on day 3 pi. Viral titers were significantly higher in aged ferrets when compared to young animals on day 5 pi (*p<*0.001) (Fig. 4b). Although differences in viral titers were observed between young and aged ferrets inoculated with 10^2^ and 10^3^ PFU at limited time points, these differences were more evident between the age groups in animals inoculated with the highest SARS-CoV-2 dose (10^6^ PFU). In these groups, infectious virus was recovered from OPS and NS (Fig. 3e, f). While young ferrets shed between 1.0 to 2.0 log10TCID_50_.mL^-1^ in OPS secretions, aged animals shed up to 3.8 log10TCID_50_.mL^-1^ of virus (days 3-5 pi, *p*<0.0001; and day 7 pi, *p<*0.001; Fig. 4c). Viral titers were above the limit of quantification (1.0 log10TCID_50_.mL^-1^) in 4/4 aged ferrets from day 1-7 pi, however, in young animals, viral titers were only detected in 2/4 on days 1 and 3 pi, and 1/4 on day 5 pi (Fig. 4c). These results indicate that SARS-CoV-2 replication was more efficient in the upper respiratory tract of aged animals. Thus, suggesting that aged ferrets are more susceptible to SARS-CoV-2 infection.

**Fig. 4.**
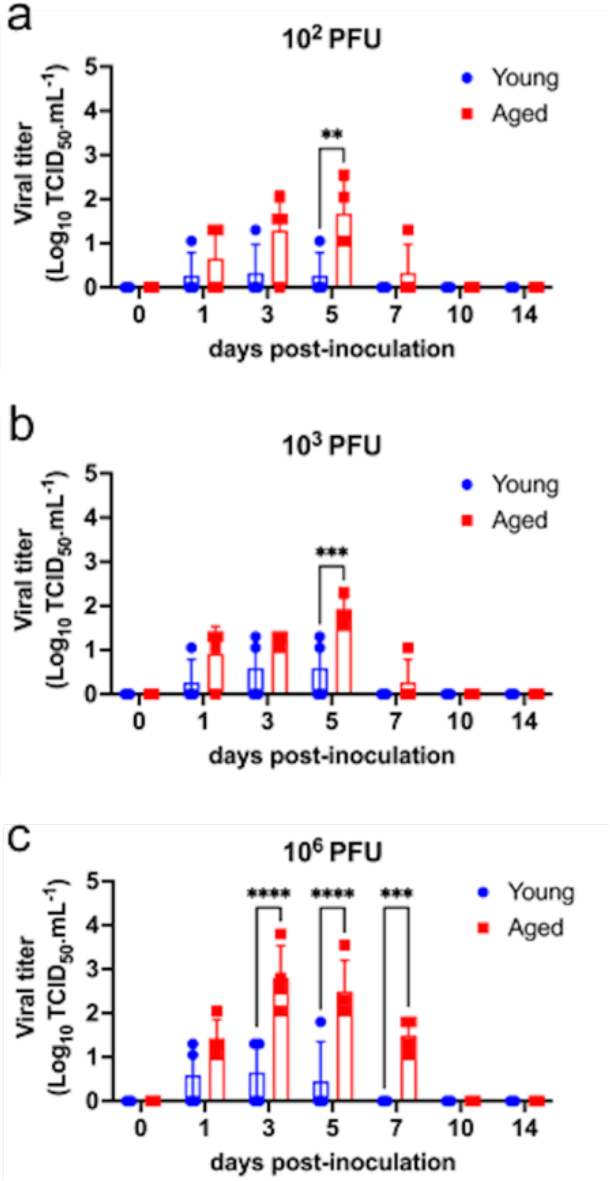
Infectious SARS-CoV-2 loads in oropharyngeal secretion. Oropharyngeal swab samples (OPS) that tested positive by rRT-PCR were subjected to virus quantitation. Viral titers in young and aged ferrets inoculated with 10^2^ PFU- (**a**), 10^3^ PFU- (**b**), or 10^6^ PFU of SARS-CoV-2 (**c**). Virus titers were determined using endpoint dilutions and the Spearman and Karber’s method and expressed as log10 TCID_50_.ml^-1^. **** = *p*<0.0001; *** = *p*<0.001; ** = *p*<0.002 * = *p*<0.02.

### Viral load in tissues

Viral RNA load and tissue distribution of SARS-CoV-2 were assessed on day 14 pi. Nasal turbinate, soft palate/tonsil, left and right lungs (cranial and caudal lobes) were collected and processed by rRT-PCR. SARS-CoV-2 RNA was only detected in nasal turbinate and soft palate/tonsil from young and aged ferrets inoculated with 10^6^ PFU (Fig. 5). Viral RNA was detected with high Ct value in the nasal turbinate of 3/4, and soft palate/tonsil 2/4 of young ferrets, while in aged animals, the nasal turbinate of 4/4 and the soft palate/tonsil of 3/4 animals were rRT-PCR positive. While the tissues were positive for viral RNA no infectious virus was recovered after three blind passages in cell culture.

**Fig. 5.**
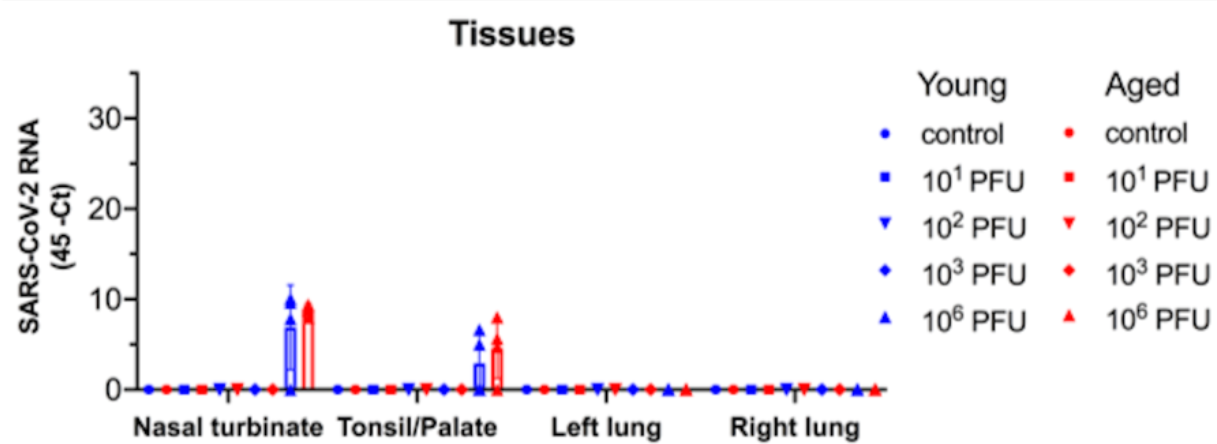
Viral RNA in tissues following SARS-CoV-2 inoculation. Viral RNA load and tissue distribution of SARS-CoV-2 were assessed on day 14 post-inoculation (pi) in nasal turbinate, soft palate/tonsil, left and right lungs (cranial and caudal lobes). Results of rRT-PCR for all ten groups doses, including young and age ferrets, are shown.

### Neutralizing antibody responses to SARS-CoV-2

The serological response to SARS-CoV-2 was assessed by virus neutralization (VN) assay. Serum samples collected on days 0, 3, 7, and 14 pi were used to determine neutralizing antibody (NA) responses following SARS-CoV-2 infection. All control (mock-inoculated) animals and those inoculated with 10^1^ PFU remained seronegative until the end of the experiment on day 14 pi (Fig. 6a). In the groups inoculated with 10^2^ PFU of SARS-CoV-2, 2/4 young and 4/4 aged ferrets seroconverted and presented NA titers on day 14 pi (Fig. 6b, *p*<0.02). All animals in the 10^3^ PFU group, regardless of age, seroconverted by day 14 pi and no differences in NA titers between age groups were detected (Fig. 6c). Among the animals inoculated with the highest dose (10^6^ PFU), both young and aged ferrets seroconverted as early as day 7 pi, with higher antibody titers being detected on day 14 pi (Fig. 6d). Together these results confirm successful infection of 2/4 young ferrets and 4/4 aged ferrets inoculated with 10^2^ PFU of SARS-CoV-2.

**Fig. 6.**
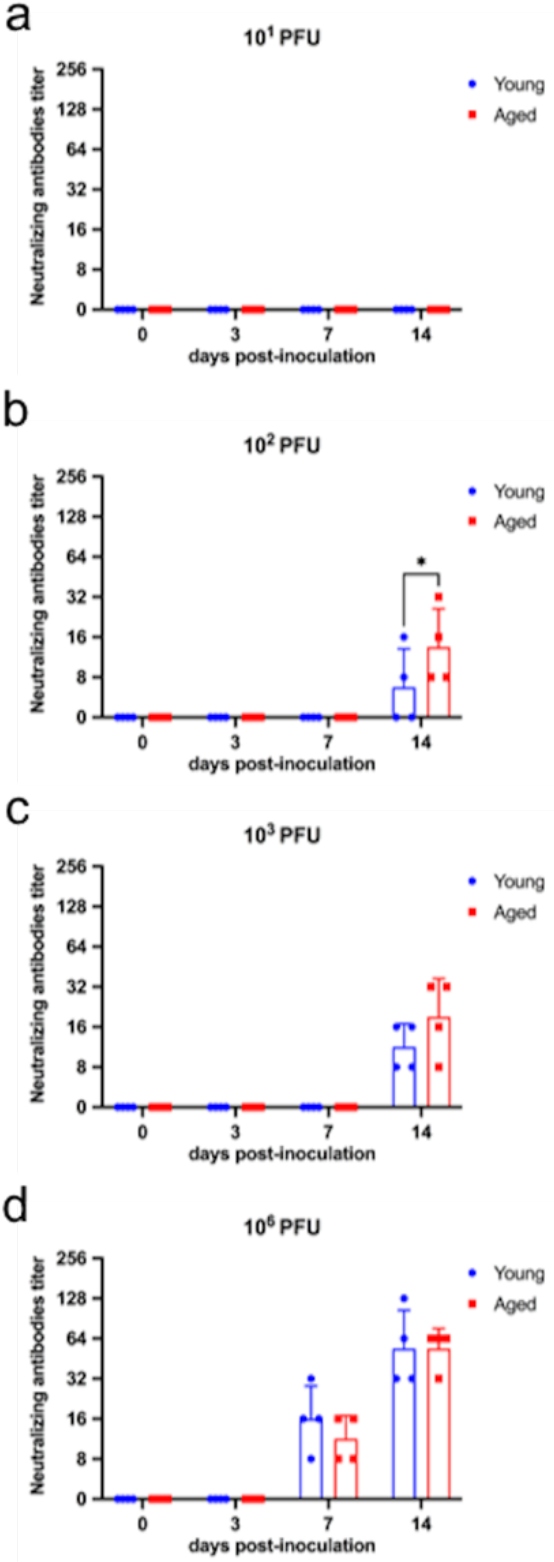
Neutralizing antibody responses to SARS-CoV-2. Serological response to SARS-CoV-2 assessed by virus neutralization assay. Serum samples collected on days 0, 3, 7, and 14 post-inoculation (pi). Neutralizing antibody responses in young and aged ferrets inoculated with 10^1^ (**a**), 10^2^ (**b**), 10^3^ (**c**), and 10^6^ PFU (**d**) of SARS-CoV-2. Neutralizing antibody titers represent the reciprocal of the highest dilution of serum that completely inhibited SARS-CoV-2 infection/replication. * = *p*<0.02.

### Median infectious dose (ID_50_) of SARS-CoV-2 in young and aged ferrets

The infectivity of SARS-CoV-2 in young and aged ferrets was assessed using three parameters: i. rRT-PCR in oropharyngeal secretion, ii. virus isolation in oropharyngeal secretion, and iii. seroconversion to SARS-CoV-2 on day 14 pi. This resulted in three binary (i.e. positive or negative) response variables for each ferret in the study. Each animal was determined to be infected when at least two of the three parameters evaluated were positive. It is important to note that rRT-PCR results alone were not used as a definitive proof of infection, shedding of infectious virus or seroconversion were also required to define an animal as infected. Given the consistency of virus shedding detected in oropharyngeal swabs (Fig. 3; Fig. 4), the frequency of young and aged ferrets shedding virus in oropharyngeal secretions that seroconverted to SARS-CoV-2 were then used to estimate the median infectious dose (ID_50_) of the virus using the three-parameter logistic dose response model.

Notably, all three parameters (rRT-PCR positive OPS, VI positive OPS and seroconversion) used to determine the infection status of young and aged ferrets provided consistent outcomes (Table 1). None of the animals in the young or aged groups shed infectious virus nor seroconverted when inoculated with 10^1^ PFU of SARS-CoV-2, indicating no infection. In animals inoculated with 10^2^ PFU, 2/4 young animals tested positive by rRT-PCR and virus isolation and the same two animals seroconverted as determined by detection of NA on day 14 pi, while all aged ferrets inoculated with 10^2^ PFU were rRT-PCR- and virus isolation positive and seroconverted to SARS-CoV-2. All young and aged animals inoculated with 10^3^ and 10^6^ PFU tested positive by rRT-PCR and infectious virus and seroconverted to SARS-CoV-2 by day 14 pi. Based on the infection status of the animals, the estimated ID_50_ for SARS-CoV-2 in aged ferrets was 31.6 PFU (∼32), while in young animals the ID_50_ was estimated at 100.1 PFU (∼100; Fig. 7).

**Fig. 7.**
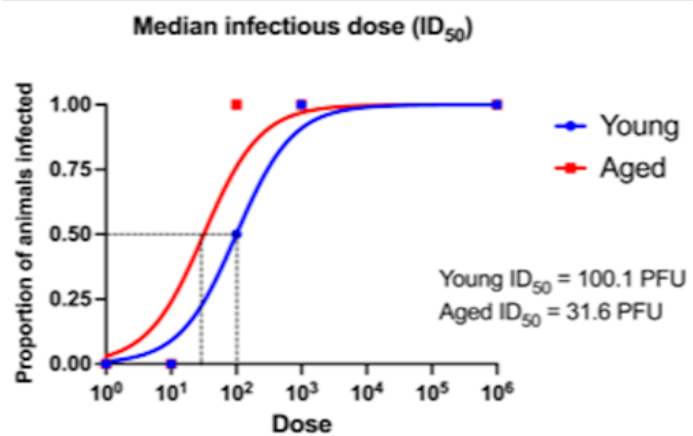
Estimated infectious dose 50 (ID_50_) of SARS-CoV-2 in young and aged ferrets. The infectivity of SARS-CoV-2 in young and aged ferrets assessed by using three parameters: i. rRT-PCR in oropharyngeal secretion, ii. virus isolation in oropharyngeal secretion, and iii. seroconversion to SARS-CoV-2 on day 14 post-inoculation. For rRT-PCR and virus isolation, an animal was considered positive if tested positive any time point throughout the 14-day experimental period. Each ferret was determined to be infected when at least two of the three parameters evaluated were positive. Median infectious dose (ID_50_) of SARS-CoV-2 was estimate using the three-parameter logistic dose response model.

**Table 1.** Median infectious dose (ID_50_) data.

| Ferret group | Young |  |  | Aged |  |  |
| --- | --- | --- | --- | --- | --- | --- |
|  | rRT-PCR | Virus isolation | Serology | rRT-PCR | Virus isolation | Serology |
| 10<sup>1</sup> PFU | 0/4 (0%) | 0/4 (0%) | 0/4 (0%) | 1/4 (25%) | 0/4 (0%) | 0/4 (0%) |
| 10<sup>2</sup> PFU | 2/4 (50%) | 2/4 (50%) | 2/4 (50%) | 4/4 (100%) | 4/4 (100%) | 4/4 (100%) |
| 10<sup>3</sup> PFU | 4/4 (100%) | 4/4 (100%) | 4/4 (100%) | 4/4 (100%) | 4/4 (100%) | 4/4 (100%) |
| 10<sup>6</sup> PFU | 4/4 (100%) | 4/4 (100%) | 4/4 (100%) | 4/4 (100%) | 4/4 (100%) | 4/4 (100%) |

### Expression of ACE2 and TMPRSS2 in the respiratory tract of young and aged ferrets

To investigate factors that could potentially contribute to susceptibility of young and aged ferrets to SARS-CoV-2, we assessed expression of ACE2 and TMPRSS2 – two critical viral entry factors – in the upper (nasal turbinates) and lower (lungs) respiratory tract of all animals in the study (young [n = 20] and aged ferrets [n = 20]) by qRT-PCR. Notably, ACE2 and TMPRSS2 mRNA levels were higher in the nasal turbinates of aged ferrets than in young animals (*p*<0.05) (Fig. 8a, b). While no differences in ACE2 and TMPRSS2 expression between age groups were observed in the lungs (Fig. 8c, d). Additionally, ACE2 expression was higher in the upper respiratory tract (URT) when compared to the lower respiratory tract (LRT) in both young and aged animals (*p*<0.05 and *p<*0.0001, respectively) (Fig. 8e, g). Expression levels of TMPRSS2, on the other hand, was higher in the LRT when compared URT in both young and aged ferrets (*p*<0.0001) (Fig. 8f, h).

**Fig. 8.**
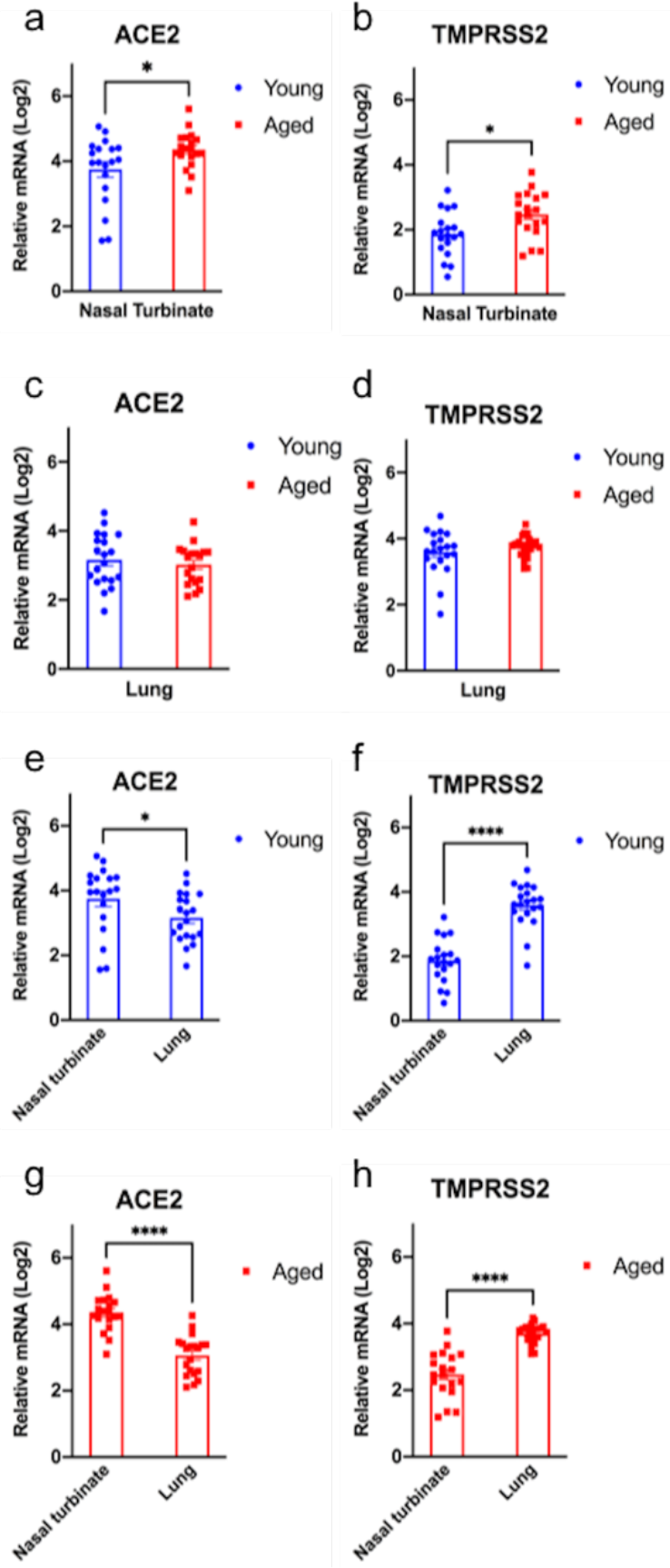
Expression of ACE2 and TMPRSS2 in the respiratory tract of young and aged ferrets. The levels of ACE2 and TMPRSS2 expression in the upper (nasal turbinates) and lower (lungs) respiratory tract of all ferrets in the study were assessed by qRT-PCR. Expression levels of ACE2 and TMPRSS2 (**a** and **b**, respectively) in nasal turbinates of young and aged ferrets. Expression levels of ACE2 and TMPRSS2 (**c** and **d**, respectively) in the lungs of young and aged ferrets. Expression levels of ACE2 and TMPRSS2 in nasal turbinates and lung of young (**e** and **f**, respectively), and aged (**g** and **h**, respectively) ferrets. **** = *p*<0.0001; *** = *p*<0.001* = *p*<0.05

## Discussion

Here we compared the susceptibility of young and aged ferrets to SARS-CoV-2 and assessed the infectivity of the virus by inoculating young and aged animals with increasing viral doses. As evidenced by SARS-CoV-2 replication in the upper respiratory tract, virus shedding in respiratory secretions (viral RNA and infectious virus) and seroconversion following intranasal inoculation, our study shows that aged ferrets are more susceptible than young ferrets to SARS-CoV-2 infection.

Intranasal inoculation of 10^1^ PFU of SARS-CoV-2 did not result in productive infection in young nor in aged ferrets. While all aged animals were successfully infected after inoculation with 10^2^ PFU/animal, only 2 of 4 young ferrets were infected with this viral dose. Importantly, the infection rates observed in young and aged animals were consistent and supported by all three criteria used do define productive infection: i. positive rRT-PCR, ii. infectious virus shedding in respiratory secretions, and iii. seroconversion (Table 1). These results suggest that the infectious dose of SARS-CoV-2 required to infect aged ferrets is lower than the dose required to infect young animals. Indeed, the ID_50_ estimated using the three-parameter logistic dose response model in aged animals was ∼32 PFU, while in young animals it was ∼100 PFU (∼3X higher). Similarly, a recent study conducted with a French SARS-CoV-2 isolate (UCN19) demonstrated successful infection of 10-month old ferrets with 2 x 10^3^ PFU of the virus^23^. Interestingly, when younger ferrets (7-month old) and a lower dose (5 x 10^2^ PFU per animal) were used, only 1 of 6 inoculated ferrets was infected after intranasal inoculation with of a SARS-CoV-2 isolate from Australia (Victoria/1/2020)^24^. Despite inherent experimental differences, observations from these earlier studies are consistent with the results presented here demonstrating a higher susceptibility of older ferrets to SARS-CoV-2. It is important to note, however, that both UCN19 and Victoria/1/2020 isolates belong to early SARS-CoV-2 lineages (lineage A) that do not contain the Spike mutation D614G, which is present in the isolate NYI67-20 (lineage B.1) used in our study. This is relevant as several studies have shown that the S D614G mutation is associated with increased infectivity and transmission of SARS-CoV-2 in humans and animal models^25–29^.

Differences in viral RNA load between age groups inoculated with same viral doses were observed mainly on days 7 and 10 pi, when aged ferrets remained positive, while viral RNA was no longer detected in most young animals. The highest viral loads were detected on OPS samples when compared to NS and RS. Higher viral loads in OPS or nasopharyngeal swab (NPS) samples have also been described in humans when compared to sputum or anterior nares samples (ANS)^30–33^. Most importantly, shedding of infectious virus in aged ferrets inoculated with SARS-CoV-2 was prolonged when compared to young animals and the viral titers detected in this age group were higher than those detected in young animals.

Infectious virus was isolated from aged ferrets with a higher frequency and for prolonged time when compared to young animals. These differences were more pronounced in animals inoculated with the highest viral dose (10^6^ PFU), from which infectious virus was isolated from all four aged ferrets from day 1 to 7 pi. Additionally, the viral titers were significantly higher in aged animals when compared to young ferrets on day 5 pi in the 10^2^ PFU (*p<*0.002) and 10^3^ PFU (*p<*0.001) groups, and on days 3, 5, and 7 pi in the 10^6^ PFU group (*p<*0.001). Together these results demonstrate that SARS-CoV-2 replicates more efficiently in aged ferrets when compared to young animals. Regardless of the viral dose, after day 7 pi, no infectious virus was isolated from any animal inoculated with the different viral doses, despite detection of viral RNA by rRT-PCR up to day 10-14 pi. These results corroborate virus shedding patterns observed in humans, in which the infectious period was shown to last 7-to-10 days following infection^34^. Importantly, in humans a decrease in SARS-CoV-2 infectivity parallels increased levels of neutralizing antibodies in serum^8, 34^. This was also observed here ferrets, which suggests that antibody responses could play a role in viral clearance. Innate immune responses at the site of virus replication, however, may also play a role and contribute to control the infection in the respiratory tract.

Our findings showing higher susceptibility of aged ferrets to SARS-CoV-2 infection in the present study mirror clinical observations in humans, which point to increased susceptibility and higher levels of virus replication in the respiratory tract of older people when compared to young children^12, 14^. Additionally, results presented here corroborate findings of a recent study by Kim and collaborators^35^, who showed age-related differences in viral load in the respiratory tract, and lung histopathology in ferrets inoculated with SARS-CoV-2. This study also showed that expression levels of genes related to the interferon (IFN) pathway, activated T cells, and macrophage responses were increased in older ferrets following SARS-CoV-2 infection^35^. These changes are likely due to enhanced immune responses following higher viral replication in aged animals. Based on our ID_50_ estimates indicating that the infectious dose of SARS-CoV-2 is ∼3X higher in young ferrets (∼100 PFU) when compared to aged animals (∼32 PFU), we hypothesized that differential expression of key SARS-CoV-2 entry factors such as the ACE2 receptor and the TMPRSS2 protease could underlie age-related differences in their susceptibility to SARS-CoV-2.

Notably, we showed that expression of both ACE2 and TMPRSS2 were lower in nasal turbinates (primary site of SARS-CoV-2 replication in the URT) of young ferrets when compared to expression levels in aged animals. Additionally, expression of ACE2 was higher in the URT when compared to the lung (LRT). These observations corroborate findings in humans^20^ and suggest that differences in expression of ACE2 and TMPRSS2 in the respiratory tract may contribute to age-related susceptibility to SARS-CoV-2. The spectrum of factors that can contribute to SARS-CoV-2 susceptibility, however, is broad and further studies are needed to assess the function and involvement of other entry and immune response-related factors on SARS-CoV-2 infection and replication in different age groups. The young and aged ferret model developed here provide an excellent platform to investigate age related differences in susceptibility to SARS-CoV-2 infection and replication and the host and viral factors that play a role in these dynamic interactions. This model may be particularly useful to dissect the mechanisms/functions emerging in several of the SARS-CoV-2 variants (e.g. P.1, B.1.617.2), which seem to have an enhanced ability to infect younger age individuals.

In summary, here we demonstrated that age affects susceptibility of ferrets to SARS-CoV-2, with aged animals being more likely to get infected when exposed to lower infectious dose of the virus when compared to young animals. Additionally, SARS-CoV-2 replication in the URT and shedding in respiratory secretions is enhanced in aged ferrets when compared to young animals. We also showed that similar to what has been described in humans^20^, aged ferrets express higher levels of ACE2 and TMPRSS2 – two key factors determining virus entry into cells – in the URT (Fig. 9). Together these results suggest that the higher infectivity and enhanced ability of SARS-CoV-2 to replicate in aged individuals is associated – at least in part – with expression levels of ACE2 and TMPRSS2 at the sites of virus entry.

**Fig. 9.**
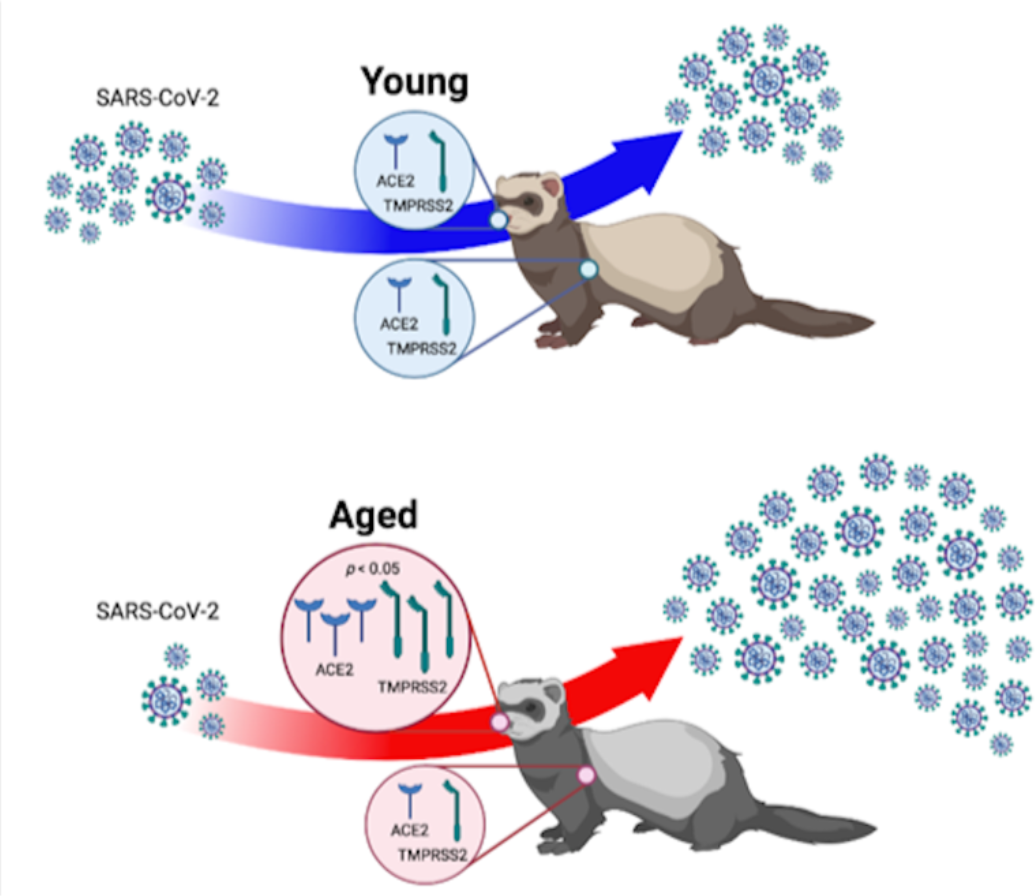
Age-related differential susceptibility to SARS-CoV-2 infection. Aged ferrets were more likely to get infected when exposed to lower infectious dose of the virus when compared to young animals. SARS-CoV-2 replication in the upper respiratory tract and shedding in respiratory secretions is enhanced in aged ferrets when compared to young animals. Notably, aged ferrets express higher levels of ACE2 and TMPRSS2 in the upper respiratory tract. Together these results suggest that the higher infectivity and enhanced ability of SARS-CoV-2 to replicate in aged individuals is associated with expression levels of two of the molecules that are critical for SARS-CoV-2 infection and host cell entry.

## Methods

### Virus and cells

Vero E6 (ATCC^®^ CRL-1586^™^), and Vero E6/TMPRSS2 (JCRB Cell Bank, JCRB1819) were cultured in Dulbecco’s modified eagle medium (DMEM), supplemented with 10% fetal bovine serum (FBS), L-glutamine (2mM), penicillin (100 U.ml^−1^), streptomycin (100 µg.ml^−1^) and gentamycin (50 µg.ml^−1^). The cell cultures were maintained at 37 °C with 5% CO_2_. The SARS-CoV-2 isolate NYI67-20 (B.1 lineage) was propagated in Vero E6 cells. Low passage virus stock (passage 3) were prepared, cleared by centrifugation (2000 x *g* for 15 min) and stored at -80 °C. The whole genome sequence of the virus stock was determined to confirm that no mutations occurred during passages in cell culture. The titer of virus stock was determined by calculated according to the Spearman and Karber method and expressed as plaque-forming units (PFU) per milliliter.

### Animals housing and experimental design

All animals were handled in accordance with the Animal Welfare Act and the study procedures were reviewed and approved by the Institutional Animal Care and Use Committee at the Cornell University (IACUC approval number 2020-0064). A total of forty ferrets (*Mustela putorius furo*) were obtained from a commercial breeder (Triple F, Gillett, PA, USA). Twenty young (6-month-old ferrets) and twenty aged ferrets (18 to 39-month-old [30.5±5.3 - average±SD]) were randomly allocated to experimental groups (n = 4). While the controls animals were housed at Animal Biosafety Level 1 (ABLS-1), all the virus inoculated ferrets were housed in the Biosafety Level 3 (ABSL-3) facility at the East Campus Research Facility (ECRF) at Cornell University. After 72 h of acclimation, ferrets were sedated and inoculated intranasally with 1 ml (0.5 ml per nostril) of a virus suspension containing 10^1^, 10^2^,10^3^, or 10^6^ PFU of SARS-CoV-2 (Groups 2, 3, 4 and 5, respectively [n = 4/group]). Ferrets of each age maintained as mock-inoculated control (Group 1, n = 4) were mock-inoculated with Vero E6 cell culture medium supernatant. All ferrets were maintained individually in Horsfall HEPA-filtered cages, connected to the ABSL-3’s exhaust system. Body temperatures and weight were measured in a daily base. Oropharyngeal (OPS), nasal (NS), and rectal swabs (RS) were collected under sedation (dexmedetomidine) on days 0, 1, 3, 5, 7, 10, and 14 post-inoculation (pi). Upon collection, swabs were placed in sterile tubes containing 1 ml of viral transport medium (VTM Corning^®^, Glendale, AZ, USA) and stored at -80 °C until processed for further analyses. Blood was collected under sedation (dexmedetomidine) through cranial vena cava venipuncture using a 3 ml sterile syringe and 23G x 1” needle and transferred into serum separator tubes on days 0, 3, 7, and 14 pi. The blood tubes were centrifuged at 1500 x *g* for 10 min and serum was aliquoted and stored at -20 °C until further analysis. Ferrets were humanely euthanized on day 14 pi under deep inhalatory anesthesia using isoflurane followed by heart puncture. Following necropsy, tissues, including nasal turbinate, soft palate/tonsil, left and right lung were collected and processed for rRT-PCR and virus isolation.

### Nucleic acid isolation and real-time reverse transcriptase PCR

Nucleic acid was extracted from OPS, NS, RS, and tissue samples collected at necropsy. For OPS, NS and RS samples 200 µL of cleared swab supernatant were used for nucleic acid extraction. For tissues, 0.2 g of each tissue were minced with a sterile scalpel, resuspended in 2 ml DMEM (10% w/v) and homogenized using a stomacher (one speed cycle of 60s, Stomacher^®^ 80 Biomaster). Then, 200 µL of the tissue homogenate supernatant was used for RNA extraction using the MagMax Core extraction kit (Thermo Fisher, Waltham, MA, USA) and the automated KingFisher Flex nucleic acid extractor (Thermo Fisher, Waltham, MA, USA) following the manufacturer’s recommendations. The real-time reverse transcriptase PCR (rRT-PCR) was performed using the EZ-SARS-CoV-2 Real-Time RT-PCR assay (Tetracore Inc., Rockville, MD, USA). An internal inhibition control was included in all reactions. Positive and negative amplification controls were run side-by-side with test samples.

### Virus isolation and titrations

All OPS, NS, and RS and tissue samples that tested positive for SARS-CoV-2 by rRT-PCR were subjected to virus isolation under Biosafety Level 3 (BSL-3) conditions. Twenty-four well plates were seeded with ∼75,000 Vero E6/TMPRSS2 cells per well 24 h prior to sample inoculation. Cells were rinsed with phosphate buffered saline (PBS) (Corning^®^, Glendale, AZ, USA), inoculated with 150 µl of each sample and the inoculum adsorbed for 1 h at 37 °C with 5% CO_2_. Mock-inoculated cells were used as negative controls. After adsorption, replacement cell culture media supplemented with FBS as described above was added, and cells were incubated at 37 °C with 5% CO_2_ and monitored daily for cytopathic effect (CPE) for 3 days. SARS-CoV-2 replication in CPE-positive cultures was confirmed with an immunofluorescence assay (IFA) as previously described ^36, 37^. Cell cultures with no CPE were frozen, thawed, and subjected to two additional blind passages/inoculations in Vero E6/TMPRSS2 cell cultures. At the end of the third passage, the cells cultures were subjected to IFA ^36, 37^. OPS were subjected to end point titrations. For this, the original sample was subjected to limiting dilutions and inoculated into Vero E6/TMPRSS2 cells cultures in 96-well plates. At 48 h post-inoculation, cells were fixed with 3.7% formaldehyde for 30 min at room temperature (rt), permeabilized with 0.2% Triton X-100 for 10 min at rt (in Phosphate Buffered Saline [PBS]) and subjected to an immunofluorescence assay (IFA) using a rabbit polyclonal antibody (pAb) specific for the SARS-CoV-2 nucleoprotein (N) (produced in Dr. Diel’s laboratory) followed by incubation with a goat anti-rabbit IgG (goat anti-rabbit IgG, DyLight^®^594 Conjugate, Immunoreagent Inc.). Virus titers were determined on each time point using end-point dilutions and the Spearman and Karber’s method and expressed as TCID_50_.ml^−1^.

### Serology

Neutralizing antibody responses to SARS-CoV-2 were assessed by virus neutralization (VN) assay performed under BSL-3 laboratory conditions. Twofold serial dilutions (1:8 to 1:1,024) of serum samples were incubated with 100 - 200 TCID_50_ of SARS-CoV-2 isolate NYI67-20 for 1 h at 37 °C. Following incubation of serum and virus, 50 µl of a cell suspension of Vero E6 cells was added to each well of a 96-well plate and incubated for 48 h at 37 °C with 5% CO_2_. The cells were fixed, permeabilized and subjected to IFA as described above. Unbound antibodies were washed from cell cultures by rinsing the cells PBS, and virus infectivity was assessed under a fluorescence microscope. Neutralizing antibody titers were expressed as the reciprocal of the highest dilution of serum that completely inhibited SARS-CoV-2 infection/replication. Fetal bovine serum (FBS) and positive and negative serum samples from white-tailed-deer fawns ^36^ were used as controls.

### Expression level of *ACE2* and *TMPRSS2* genes in the respiratory tract

RNA samples extracted from nasal turbinate and lungs (described above) were treated with DNA-*free*^TM^ Kit DNase Treatment & Removal (Invitrogen, Carlsbad, CA, USA) according the manufacturer’s instructions. Thus, the amount of RNA in each sample was measured using Qubit^TM^ RNA BR Assay Kit (Invitrogen, Carlsbad, CA, USA) and then all the samples were diluted in ddH2O in order to obtain a concentration of 5 ng/µl. Standard curves were prepared from a pool of RNA of all samples (2-fold dilutions). Custom primers and probe were designed to angiotensin-converting enzyme 2 (ACE2), transmembrane serine protease 2 (TMPRSS2) and glyceraldehyde 3-phosphate dehydrogenase (GAPDH) of ferrets using PrimerQuest Tool from Integrated DNA Technologies website (https://www.idtdna.com/pages). The primers and probe sequence for Ferret ACE2 were 5’-GATGTGAGGGTGAGCGATTT-3′, 5′-GGGACTTCCTGATAGCTTCTTC3′ and /56-FAM/TGACATCAT/ZEN/TCCCAGAGCTGACGT/3IABkFQ/ based on *Mustela putorius furo* angiotensin converting enzyme 2 (ACE2) GenBank accession number NM_001310190.1; for Ferret TMPRSS2 were 5’-CGGTGTTTACGGACTGGATTTA-3′, 5′-GGTGCCCAGAGAATGAAGAA-3′ and /56-FAM/ACAGCTAAT/ZEN/CCATGTGCCCTGTGT/3IABkFQ/ based on *Mustela putorius furo* serine transmembrane protease 2 (TMPRSS2) GenBank accession number NM_001386127.1; and for Ferret GAPDH were 5’-GATGCTGGTGCTGAGTATGT-3′, 5’-CAGAAGGAGCAGAGATGATGAC-3′ and /56-FAM/TTCACCACC/ZEN/ATGGAGAAGGCTGG/3IABkFQ/ based on *Mustela putorius furo* glyceraldehyde-3-phosphate dehydrogenase (GAPDH) GenBank accession number NM_001310173.1. Real-time RT-PCR amplifications were performed in 10-µl reactions, with 3 µl of RNA, 5 µl of TaqMan® RT-PCR mix (TaqMan® RNA-to-Ct^TM^ 1-Step Kit) (Applied Biosystems, Waltham, MA, USA), 0.5 µl of TaqMan® RT-Enzyme Mix, 0.25 µl of the mixture of primers and probe (PrimeTime qPCR probe assays) (Integrated DNA Technologies Inc., Coralville, IA, USA), and 1.25 µl of ddH2O. Amplification and detection were performed under following conditions: 15 min at 48 °C for reverse transcription, 10 min at 95 °C for polymerase activation and 40 cycles of 15 s at 95 °C for denaturation and 1 min at 60 °C for annealing and extension. The measurement of gene expression was performed by using the relative quantitation method ^38^. Relative genome copy numbers were calculated based on the standard curve determined for each gene within CFX MaestroTM software (Bio-Rad, Hercules, CA, USA), and expression levels of the genes tested were normalized to the housekeeping gene GAPDH. The amount of relative mRNA detected in each sample was expressed as log2 genome copy number.

### Median infectious dose (ID_50_) calculations

Median infectious dose (ID_50_) of SARS-CoV-2 in young and aged ferrets was assessed by using three parameters: i. rRT-PCR in oropharyngeal secretion, ii. virus isolation in oropharyngeal secretion, and iii. seroconversion to SARS-CoV-2 on day 14 pi. This resulted in three binary (i.e. positive or negative) response variables for each ferret in the study. The frequency (%) of animals positive for each dose inoculated in both young and aged group was determined (Table 1). For rRT-PCR and virus isolation, an animal was considered positive if tested positive any time point throughout the 14-day experimental period. This data was used to calculate median infectious dose (ID_50_). The three-parameter logistic model (3PL) was used to estimate ID_50_. To determine dose-response curve using 3PL model, viral dose was log transformed, bottom response value was constrained to 0 and top response value was constrained to 1. The three-parameter logistic model was implemented in GraphPad Prism software (Version 9.0.1).

### Statistical analysis

Statistical analysis was performed by 2way ANOVA followed by multiple comparisons and by unpaired t-test. Statistical analysis and data plotting were performed using the GraphPad Prism software (Version 9.0.1).

## Acknowledgements

We would like to thank the Center for Animal Resources and Education (CARE) staff, and Cornell Biosafety team for the support. This work was funded by the Office of the Vice Provost for Research, Cornell Rapid Research Response to SARS-CoV-2.

## Author contributions

M.M and D.G.D. conceived the studies and investigations. M.M, M.H.V.F, L.R.J., performed the animal procedures at ABSL-3. MM performed viral investigations and prepared figures.

M.H.V.F performed the ACE2 and TMPRSS2 expression analysis. L.R.J. performed the ID_50_ calculation. M.M. and D.G.D. wrote the manuscript. All authors critically reviewed and approved the final version of the manuscript.

## References

1. Gorbalenya, A. E. et al. The species Severe acute respiratory syndrome-related coronavirus: classifying 2019-nCoV and naming it SARS-CoV-2. Nature Microbiology (2020) doi:10.1038/s41564-020-0695-z.

2. Zhou, P. et al. A pneumonia outbreak associated with a new coronavirus of probable bat origin. Nature (2020) doi:10.1038/s41586-020-2012-7.

3. Murakami, S. et al. Detection and Characterization of Bat Sarbecovirus Phylogenetically Related to SARS-CoV-2, Japan. Emerg. Infect. Dis. (2020) doi:10.3201/eid2612.203386.

4. Lau, S. K. P. et al. Possible Bat Origin of Severe Acute Respiratory Syndrome Coronavirus 2. Emerg. Infect. Dis. (2020) doi:10.3201/eid2607.200092.

5. Wu, F. et al. A new coronavirus associated with human respiratory disease in China. Nature (2020) doi:10.1038/s41586-020-2008-3.

6. Masters, Paul s., Perlman, S. CHAPTER 28 – Coronaviridae. Fields Virol. 6th *Ed.* (2013).

7. Chen, B. et al. Overview of lethal human coronaviruses. Signal Transduct. Target. Ther. 5, (2020).

8. Cevik, M., et al. SARS-CoV-2, SARS-CoV, and MERS-CoV viral load dynamics, duration of viral shedding, and infectiousness: a systematic review and meta-analysis. The Lancet Microbe 2, e13–e22 (2021).

9. Wu, Z. & McGoogan, J. M. Characteristics of and Important Lessons From the Coronavirus Disease 2019 (COVID-19) Outbreak in China. JAMA (2020) doi:10.1001/jama.2020.2648.

10. Bialek, S. et al. Severe Outcomes Among Patients with Coronavirus Disease 2019 (COVID-19) — United States, February 12–March 16, 2020. MMWR. Morb. Mortal. Wkly. Rep. (2020) doi:10.15585/mmwr.mm6912e2.

11. Nachtigall, I. et al. Clinical course and factors associated with outcomes among 1904 patients hospitalized with COVID-19 in Germany: an observational study. Clin. Microbiol. Infect. (2020) doi:10.1016/j.cmi.2020.08.011.

12. Euser, S. et al. SARS-CoV-2 viral load distribution reveals that viral loads increase with age: a retrospective cross-sectional cohort study. medRxiv 2021.01.15.21249691 (2021) doi:10.1101/2021.01.15.21249691.

13. O’Driscoll, M. et al. Age-specific mortality and immunity patterns of SARS-CoV-2. Nature 590, 140–145 (2021).

14. Magleby, R. et al. Impact of Severe Acute Respiratory Syndrome Coronavirus 2 Viral Load on Risk of Intubation and Mortality Among Hospitalized Patients With Coronavirus Disease 2019. Clin. Infect. Dis. (2020) doi:10.1093/cid/ciaa851.

15. Letko, M., Marzi, A. & Munster, V. Functional assessment of cell entry and receptor usage for SARS-CoV-2 and other lineage B betacoronaviruses. Nat. Microbiol. (2020) doi:10.1038/s41564-020-0688-y.

16. Hoffmann, M. et al. SARS-CoV-2 Cell Entry Depends on ACE2 and TMPRSS2 and Is Blocked by a Clinically Proven Protease Inhibitor. Cell 181, (2020).

17. Chen, M. et al. Elevated ACE-2 expression in the olfactory neuroepithelium: Implications for anosmia and upper respiratory SARS-CoV-2 entry and replication. Eur. Respir. J. 56, 19–22 (2020).

18. Hamming, I. et al. Tissue distribution of ACE2 protein, the functional receptor for SARS coronavirus. A first step in understanding SARS pathogenesis. J. Pathol. 203, 631–637 (2004).

19. Sungnak, W. et al. SARS-CoV-2 entry factors are highly expressed in nasal epithelial cells together with innate immune genes. Nat. Med. 26, 681–687 (2020).

20. Singh, M., Bansal, V. & Feschotte, C. A Single-Cell RNA Expression Map of Human Coronavirus Entry Factors. Cell Rep. 32, 108175 (2020).

21. Bunyavanich, S., Do, A. & Vicencio, A. Nasal Gene Expression of Angiotensin-Converting Enzyme 2 in Children and Adults. JAMA 323, 2427 (2020).

22. Larrat, S. & Medicine, D. E. S. Z. Ferret Behavior Medicine Ferret Mustela furo Behavior Physiology Aggression. 24, 37–51 (2021).

23. Monchatre-Leroy, E. et al. Hamster and ferret experimental infection with intranasal low dose of a single strain of SARS-CoV-2. J. Gen. Virol. 102, (2021).

24. Ryan, K. et al. Dose-dependent response to infection with SARS-CoV-2 in the ferret model: evidence of protection to re-challenge. 2, 1–39 (2020).

25. Hou, Y. J. et al. SARS-CoV-2 D614G variant exhibits efficient replication ex vivo and transmission in vivo. Science *(80-.).* (2020) doi:10.1126/science.abe8499.

26. Zhou, B. et al. SARS-CoV-2 spike D614G change enhances replication and transmission. Nature 592, 122–127 (2021).

27. Plante, J. A. et al. Spike mutation D614G alters SARS-CoV-2 fitness. Nature 592, 116–121 (2021).

28. Korber, B. et al. Tracking Changes in SARS-CoV-2 Spike: Evidence that D614G Increases Infectivity of the COVID-19 Virus. Cell (2020) doi:10.1016/j.cell.2020.06.043.

29. Volz, E. et al. Evaluating the effects of SARS-CoV-2 Spike mutation D614G on transmissibility and pathogenicity. Cell 64–75 (2021) doi:10.1101/2020.07.31.20166082.

30. Hanson, K. E. et al. Self-collected anterior nasal and saliva specimens versus health care worker-collected nasopharyngeal swabs for the molecular detection of SARS-CoV-2. J. Clin. Microbiol. 58, 1–5 (2020).

31. Zhou, Y. & OLeary, T. J. Relative sensitivity of anterior nares and nasopharyngeal swabs for initial detection of SARS-CoV-2 in ambulatory patients: Rapid review and meta-Analysis. PLoS One 16, 1–12 (2021).

32. Wang, X. et al. Comparison of nasopharyngeal and oropharyngeal swabs for SARS-CoV-2 detection in 353 patients received tests with both specimens simultaneously. Int. J. Infect. Dis. 94, 107–109 (2020).

33. Lee, R. A. et al. Performance of Saliva, Oropharyngeal Swabs, and Nasal Swabs.

34. Jones, T. C. et al. Estimating infectiousness throughout SARS-CoV-2 infection course. Science (80-.). 373, (2021).

35. Kim, Y.-I. et al. Age-dependent pathogenic characteristics of SARS-CoV-2 infection in ferrets. Research square (2021).

36. Palmer, M. V. et al. Susceptibility of white-tailed deer ( Odocoileus virginianus) to SARS-CoV-2. J. Virol. (2021) doi:10.1128/jvi.00083-21.

37. Carvallo, F. R. et al. Severe SARS-CoV-2 Infection in a Cat with Hypertrophic Cardiomyopathy. Viruses 2021, Vol. 13, Page 1510 **13**, 1510 (2021).

38. Wong, M. L. & Medrano, J. F. One-Step Versus Two-Step Real-Time PCR. 39, 75–85 (2005).

